# Phytochemical screening, GC-MS profiling of invasive cocklebur (*Xanthium strumarium*)-insect-pathogen interaction and simulated volatile chemical signaling at Northeast China

**DOI:** 10.1101/2020.01.17.910240

**Authors:** Mazher Farid Iqbal, Yu Long Feng

**Affiliations:** Liaoning Key Laboratory for Biological Invasions and Global Changes, College of Bioscience and Biotechnology, Shenyang Agricultural University, Shenyang, 110866, Liaoning Province, China

**Keywords:** Biological control agents, *gall forming insects*, *Puucinia xanthii*, damage quantification, Temperature, Longitudes, Rust infected leave chemistry

## Abstract

Invasive cocklebur (*Xanthium strumarium*) is characterized by its excellent genetic and ecological plasticity, ability to spread in agriculture crops. There is a dire need to locate useful management strategies to control this invasive weed at diversified latitudinal gradients. In ecology, there is weak evidence that the damage caused by the natural enemy varies with latitudes. Therefore, we investigated this evidence with the help of transect quadratic ecological sampling method which was conducted randomly at ten different sites of Northeast China. Overall, significantly high infestation of gall-forming insect (*Epiblema strenuana*) was observed on *Xanthium* leaves (21.16%) at 41.51279°N, followed by 40.2241°N latitude. Similarly, there was a significantly high abundance of *Epiblema* infestation (7.3) with high damage (3.88%) at 41.51279°N and 40.12749°N latitude. Likewise, the fungal abundance (5.6) of rust i.e. *Puccinia xanthii* (presenting 16.23% attack) was dominated significantly at 41.51279°N. Hence, high pathogen infection rate (8.97%) was detected at 40.2241°N. On the other hand growth parameters, i.e. plant height (cm), stem diameter (mm) vary with latitude and longitudinal trends. In our experiment, of plant natural enemy interaction provides the evidence-based indication the *Epiblema* abundance, was diversified at 41.51279°N, and *P. xanthii* infection was most frequent at 40.22411°N latitudes. This study provides an evidence-based indication that natural enemy pressure varies with latitude, however this investigations gave valuable information that insect and phytopathological fungus having biological control potential against *Xanthium strumarium* invasive weed. Secondly, phytochemical qualitative and chemical signaling through Gas Chromatography-mass spectrometry (GC-MS) executed the presence of flavonoids, phenols, saponins, alkaloids, terpenoids, nitrogen (N), sulpher (S), silicon (Si) containing compounds in both treated and controlled leaves that defend against *Puccinia xanthii*. Fascinatingly, all *X. strumarium* populations collected from different latitudes possess similar compositions. In interaction mechanism, plant known to omit volatile organic compounds in response to attack of natural herbivores. The leave chemical profiling suggested that the influence of fungus attack on invasive weed brought different changes in chemical infrastructure of leave and these chemicals also play a vital role in the food web. After attack of these biological control agents, plants exhibits passionate compound reprogramming within the leaf naturally that act upon in defense systems.

**Author summary:**

- The study was conducted to observe the environmental impact on the trend of insect, invasive weed and pathogens.
- There was a significant dominance of gall-forming insect on invasive *Xanthium* weed at all locations.
- *Puccinia xanthii* infected more than 16% plantation
- Plant growth had significant variation at various longitudes and latitudes.
- The abundance of insect was positively linked with different environmental factors and *Xanthium* plant.
- The results of GC-MS suggested that *Puccinia xanthii* infected (treatment) leaves covered maximum area (%) compared to control treatments due to breakdown of the chemical compounds that proved our hypotheses that volatile organic compounds altered infrastructures of the leave chemistry that led to activeness of plant defensive chemicals resulted invasion success.

## 1. Introduction

Invasive alien species (IAS) are the plants introduced or spread from outside their natural allocations negatively affect biological biodiversity, reduce native species abundance and cause economic loss [1]. The phenomenon of plants-natural enemies interactions are complex and poorly understood [2]. This weed cannot be controlled easily without herbicides; however, studies suggested that glyphosate herbicide may be effective management to control this weed. But problem is that huge usage of chemical herbicides definitely liberates fumes to the natural ecosystem resulted to cause health hazards to the human beings and naturally grown populations in ecosystem functioning. So in this scenario biological control is best option to overcome the invasiveness of the weeds in the present circumstances.

In the recent era, the researchers have explored the changes in plant-herbivore interactions with latitudinal gradients. The latitudinal trends in plant-natural enemy interactions showed that low latitudes plant species were less palatable to natural enemies than conspecifics from higher latitudes. The natural enemies’ pressure and their interactions with plants were comparable with different latitudes. For several decades, significant studies have focused on different levels of latitudes with reference to plant natural enemy connections. In this interaction mechanism, natural enemy attack increases in lower latitudes due to the expanded season than higher latitude [3-10].

The meta-analysis found that 37% of 38 studies showed significant negative latitudinal gradients in plant natural enemy interaction, while 21% reported significant positive pattern. However, 14 comparisons recorded higher herbivory at lower latitudes, while 14 studies showed a non-significant relationship. Latitudinal comparisons of herbivore preferences were reviewed; out of which 27 studies, 59% suggested that weeds from elevated latitudes are more vulnerable to herbivores. However, there was the non-significant difference in weeds species from high versus low latitudes were found in 16 evaluations. Moreover, three comparisons recorded that natural enemies consume weed species from lower latitudes. Indeed, the diversity of natural enemy on plants can be related to latitudinal gradients. Previous findings explain the reality that chewing damage played a vital role in plant herbivore interaction mechanism with latitudinal gradients along with different climatic factors [8,9,11-18].

Stem galling moth (*Epiblema strenuana*) (Lepidoptera: Tortricidae) is native to North America that was introduced to Australia from Mexico for controlling *Xanthium occidentale, Ambrosia artemisifolia* and *Parthenium hysterophorus* in 1982. *E. strenuana* is imported to China for the management of *Ambrosia artemisiifolia*, and released on a small scale in Hunan Province, China in 1990. Then, this phytophagous insect adapted there and *Ambrosia, Xanthium* and *Parthenium* species were injured by *E. strenuana* [19]. *E. strenuana* inhibits the size, abundance and pollen production of the invasive weed [20,21]. Contradictory observations were recorded that *E. strenuana* completed its developmental stages on sunflower crops, but its economic damage was considered low. Some studies reported that gall damage slowed down the process of increasing plant height up to 40% in Australia [20,22,23].

Rust (*Puccinia xanthii*) is an obligate parasite on *Xanthium* species that overwinters in the form of spores in dead plants [24] and are well known as highly specialized pathogens [25]. The disease reduces plant growth of *Xanthium* species significantly, its dark brown telia identified on leaves [26,27]. However, optimum temperature (24 °C) and relative humidity (70%) are required [26] for its infection. *P. xanthii* was recorded on a native *Xanthium* complex, which was climatically adapted to the tropical environment in Australia [28]. However, *P. xanthii* is an effective Biological Control agent of *A. trifida* and *Xanthium* species in Australia [29]. Similarly, this rust was also observed in China, causing a severe attack on *A. trifida* [30].

Plants micro organisms adopted various defensive tactics according to the simulation environmental conditions [31,32]. In pathogen damage the plant defense reactions are self defined that frighten against tissue injury and restructuring of physiological and molecular actions to defend against the attack of pathogens [33]. Plant inherent invulnerability and its development have been systematically assessed in perception of plant-pathogen interactions [34-36]. Plant cell wall is the first leading construction layer that directly approach the pathogen, after its attack the plant trigger resistance against it [37]. Plant growth regulators already present within the plants that provide suitable signals against the harassment of the pathogen resulted to provide aid to the consequent comeback against herbivores [38]. Hence the defensive system activated resulted molecular physiology, biochemistry, volatile and non volatile (VOC’s) compounds [35,39-42] of the weed plant changed [39,43].

The weed plants correspond through VOC’s signals released as a response to pathogen attack [44,45]. The response of the plant to rust is different compared to insects or non-obligate parasites [46,47] which show sequential vibrant VOC release models [48,49]. Furthermore the understandings of plant-pathogen interaction (Fig. 9) are critical not only involved in ecological point of view but also played a significant role in IPM.

The current study investigates the relationships of two natural enemies’ having inherent biological control potential against this toxic weed. The hypotheses explored that natural enemy abundance and its damage on the leaf tissues played a significant role in the ecosystem. This study aimed to examine previous conflicting evidence that natural enemy damage varies with latitude. The goal of the study was to find out the impact of the *Epiblema strenuana* and *Puccinia xanthii* attack on *X. strumarium* in terms of biotic and abiotic factors at diversified latitudes at northeast China. It is also hypothesized that natural enemies are more abundant and develop maximum pressure on invasive weed in high latitude than in low latitude, and the natural enemies’ pressure depends on the abiotic factors. Secondly, the pathogen attacks on the leave tissues may cause transformations in volatile organic compounds altered infrastructures that led to activeness of plant defensive chemicals compared to control.

## 2. Results

### 2.1. Distribution of *Epiblema strenuana* and *Puccinia xanthii* with latitudes

Maximum *Epiblema* abundance was recorded at 41.5128°N latitude at Fushun. The relationship of Fisher’s test analysis of variance regarding this insect abundance with latitude recorded a highly significant result (P = 0.005 and F = 3.11) (Fig. 3A). The nested ANOVA of insect abundance with latitude showed a highly significant result with P = 0.000 and F = 4.450. From figure 3B, significant (P<0.05) *Epiblema* damage (%) was recorded at 41.5128°N and 41.5111°N latitude followed by 40.2244°N with P = 0.029 and F = 2.31 at a confidence level of 95%. The nested ANOVA of insect damage (%) with sites showed a significant result with P = 0.030 and F = 2.409. Figure (3C) showed statistically significant (P<0.05) prominence value (PV) recorded in F = 2.48 at 41.5128°N and 41.4949°N; however the low PV recorded at 40.4334°N.

**Figure 1:**
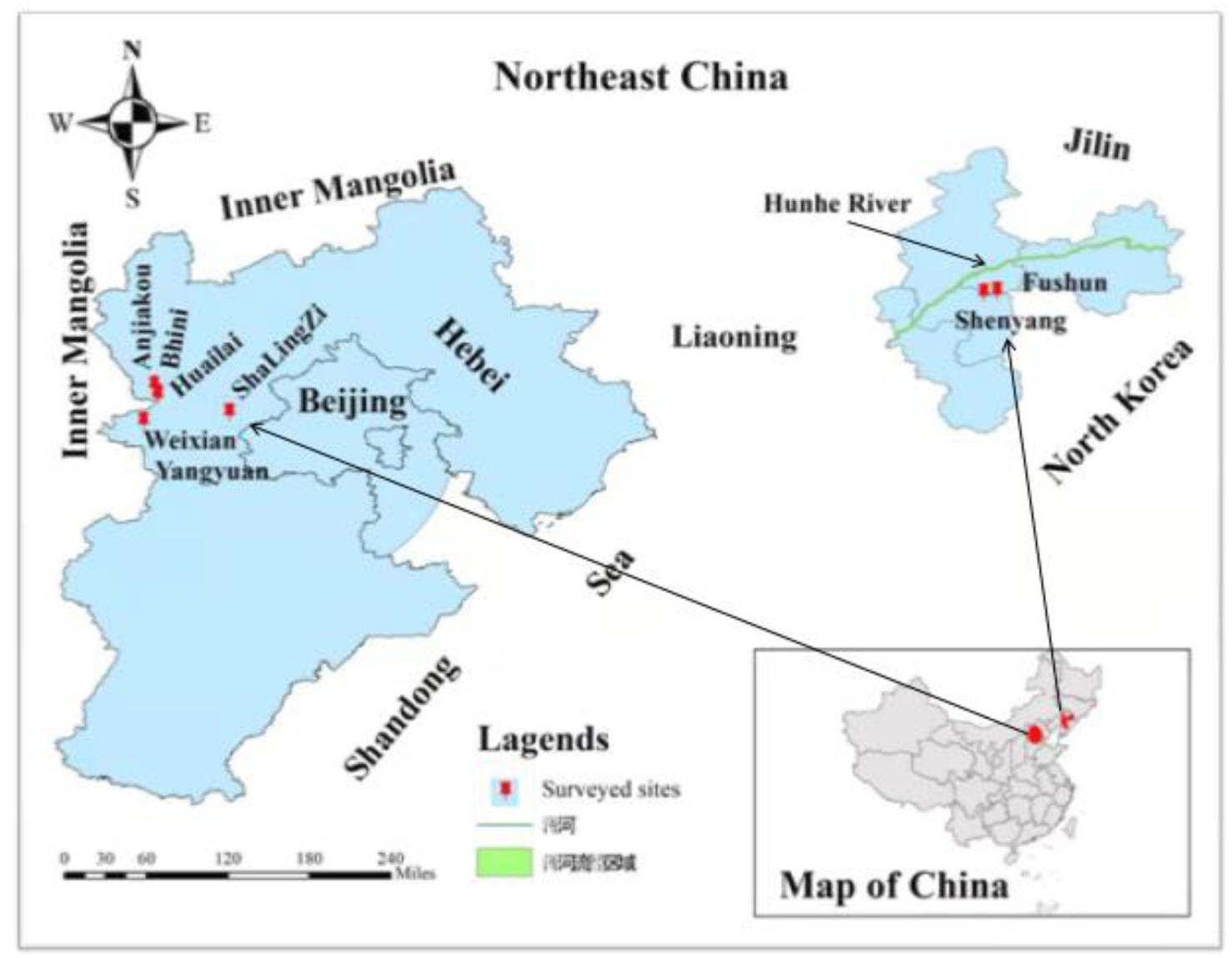
Study sites of Liaoning and Hebei Provinces in Northeast China in 2018 using ArcGIS

**Figure 2:**
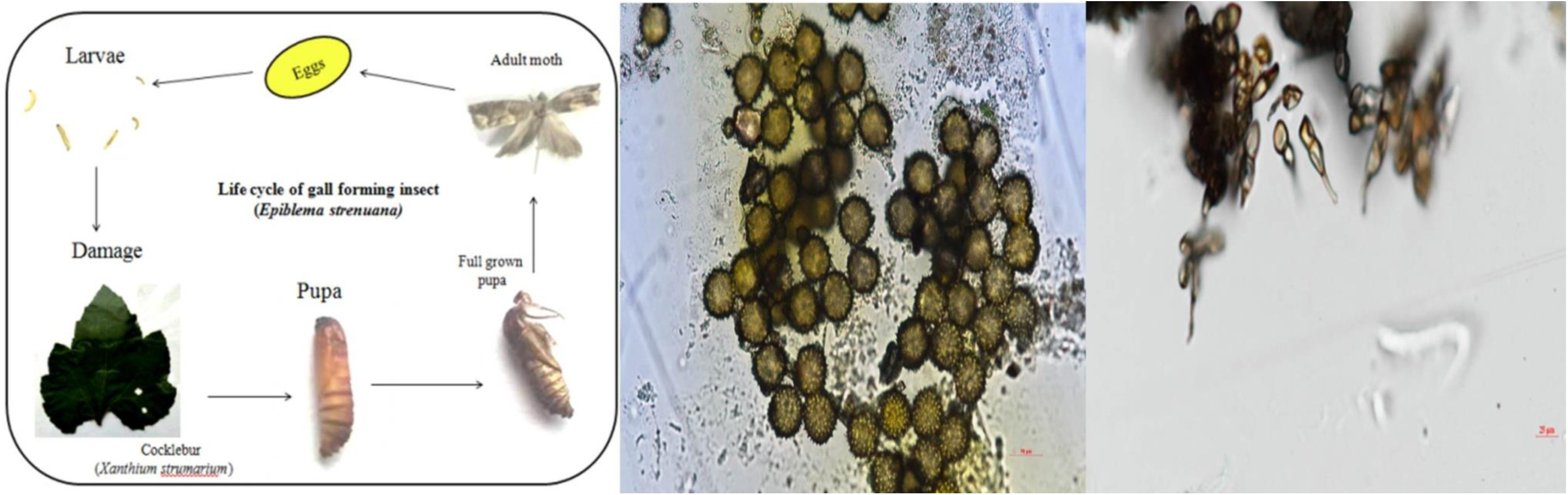
Light microscopic view of life cycle of *Epiblema strenuena* and spores of *Puccinia xanthii* on *Xanthium strumarium leaves*

**Figure 3.**
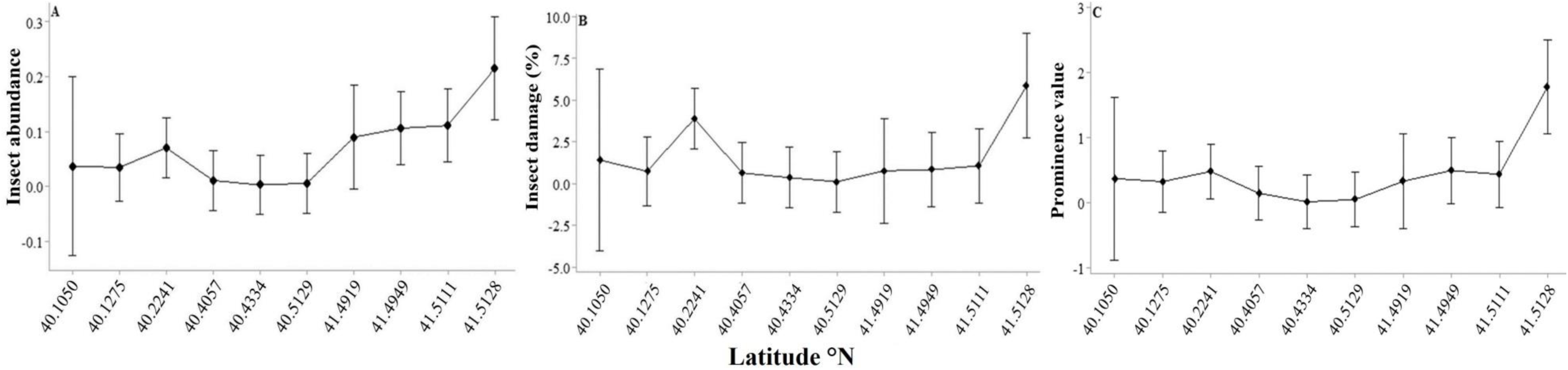
Relationship of Latitudinal patterns with insect abundance, insect damage (%) and prominence value at different latitudes

Highly significant pathogen abundance (P<0.001 and F = 11.79) was investigated at 41.49190°N, followed by 41.51111°N. The result indicated that pathogen abundance increased with an increasing trend of latitudinal gradients. Pathogen abundance was increased at 40.2241°N, but this pressure significantly increased at 41°N (Fig. 4C). The pathogen damage (%) per plant recorded maximum damage (%) on *Xanthium* leaves at 40.2241°N at Huailai County after this, its pressure decreased dramatically. Therefore, with the increasing trend of latitudinal pattern towards 41°N in the northeast area, damage pressure increased gradually. The overall *P. xanthii* pressure was maximum at a 41°N latitude with P = 0.007 and F = 2.90 (Fig. 4A). The nested ANOVA of pathogen damage (%) with the latitude showed a highly significant pattern at high latitude (P = 0.001 and F = 4.071).

**Figure 4.**
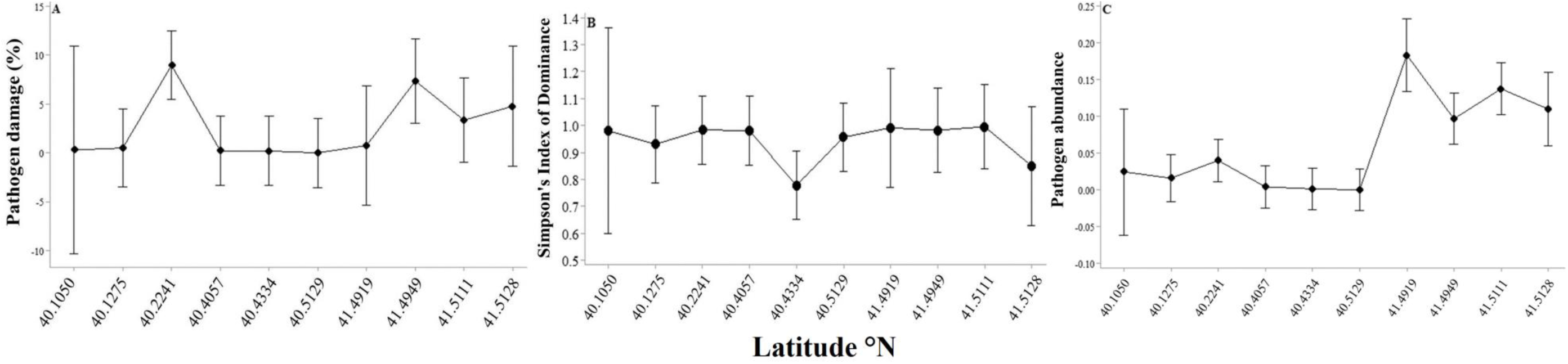
Relationship of pathogen damage (%), Simpson Index of Dominance and pathogen abundance on Xanthium strumarium with maximum annual temperature along with Latitudinal gradients

Mean July Temperature (MAT) played a vital role in the abundance and flourishing of *P. xanthii* infection. Significant abundance was investigated at 36.1 °C and 37.7 °C. The trend of pathogen abundance decreased significantly (P<0.001) with F = 22.57 (Fig. 5C) across the lower latitude sites might be due to low humidity (%). On 54.75% humidity, pathogen abundance increased gradually in northeast areas which prove the evidence that disease pressure increased with increasing level of relative humidity. The maximum Simpson Index of Dominance (SID) recorded non-significant (P>0.05) result on latitudinal gradients at lower to high (40°N to 41°N) latitudes. After 41°N, the pressure of SID decreased significantly (Fig. 4B). The results indicated that *Epiblema strunuana* and *Puccinia xanthii* were statistically significant (P<0.05) and vary across the latitudinal gradients.

**Figure 5.**
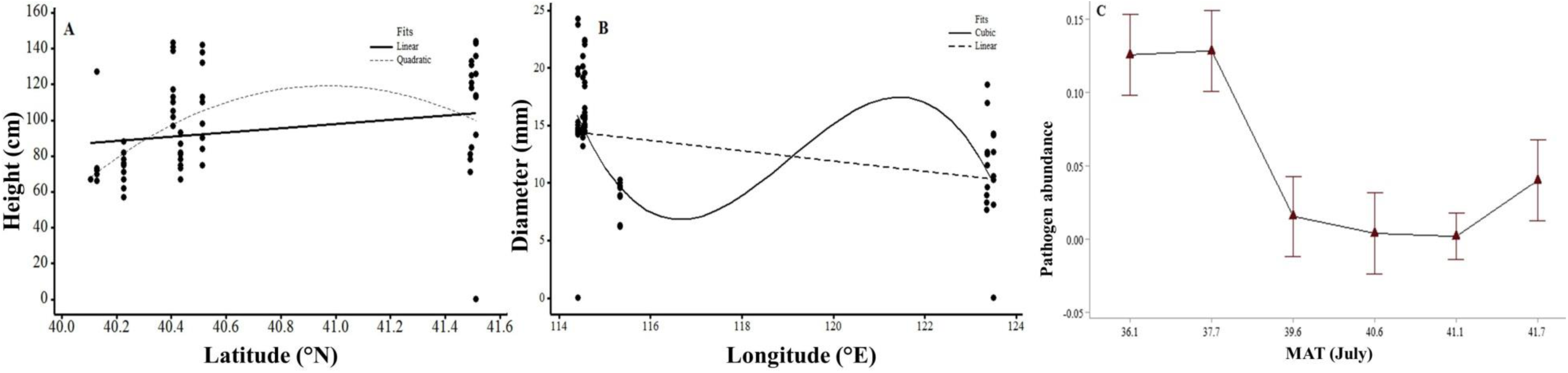
Relationship of Latitudinal, longitudinal pattern and Maximum Annual Temperature (MAT) with growth parameter and pathogen abundance on *Xanthium strumarium* invasive weed

### 2.2. Effect of latitude on growth parameters of *Xanthium strumarium*

The trend of plant height (cm) increased positively with latitude from 40°N to 41°N (Fig. 5A). The analysis of variance of plant height (cm) with nested sites showed a significant result with P = 0.02 and F = 3.660. The stem diameter (mm) of the *Xanthium* decreased significantly with increasing longitudinal trends (Fig. 5B) indicating a negative relationship. The ANOVA of diameter (mm) with nested sites showed a highly significant result (P = 0.000 and F = 4.713) and gained a maximum size at a lower longitudinal pattern.

### 2.3. Effect of *Epiblema strenuana* and *Puccinia xanthii* with abiotic factors

The graphical representation indicated that mean annual temperature (°C), relative humidity (%), mean annual rainfall (mm) played a significant role in the flourishing and spreading natural enemy pressure in the ecosystem (Figure 9). Our hypothesis is correct that insect (*Epiblema struneuna*) and pathogen (*Puccinia xanthii*) attack was recorded maximum on mean annual temperature (MAT) in July at 36°C. The potential pressure of these two natural enemies decreased significantly and correlated negatively with the increasing trends of temperature (Fig. 6A and 6B). The pathogen pressure asymptomatically reduced with the rising trend of temperature up to 42°C at 40°N latitude, which was a clear indication, that *P. xanthii* pressure decreased at lower latitude. The palatability of *Epiblema* increased with 62% to 68% relative humidity in this community (Fig. 6C and 6F) and correlated positively. The pathogen damage (%) correlated positively and showed significantly high pressure (P<0.001) with relative humidity with F = 5.87 (6F). However, the attack of the two natural enemies depends upon the mean annual precipitation (mm) and humidity level (6D and 6E). Significantly, high insect damage (%) was recorded on leaves at 50.17% relative humidity, which was decreased gradually. However, after increasing 61% RH, insect damage (%) was increased at 68% with F = 2.90 value recorded statistically (6E).

**Figure 6:**
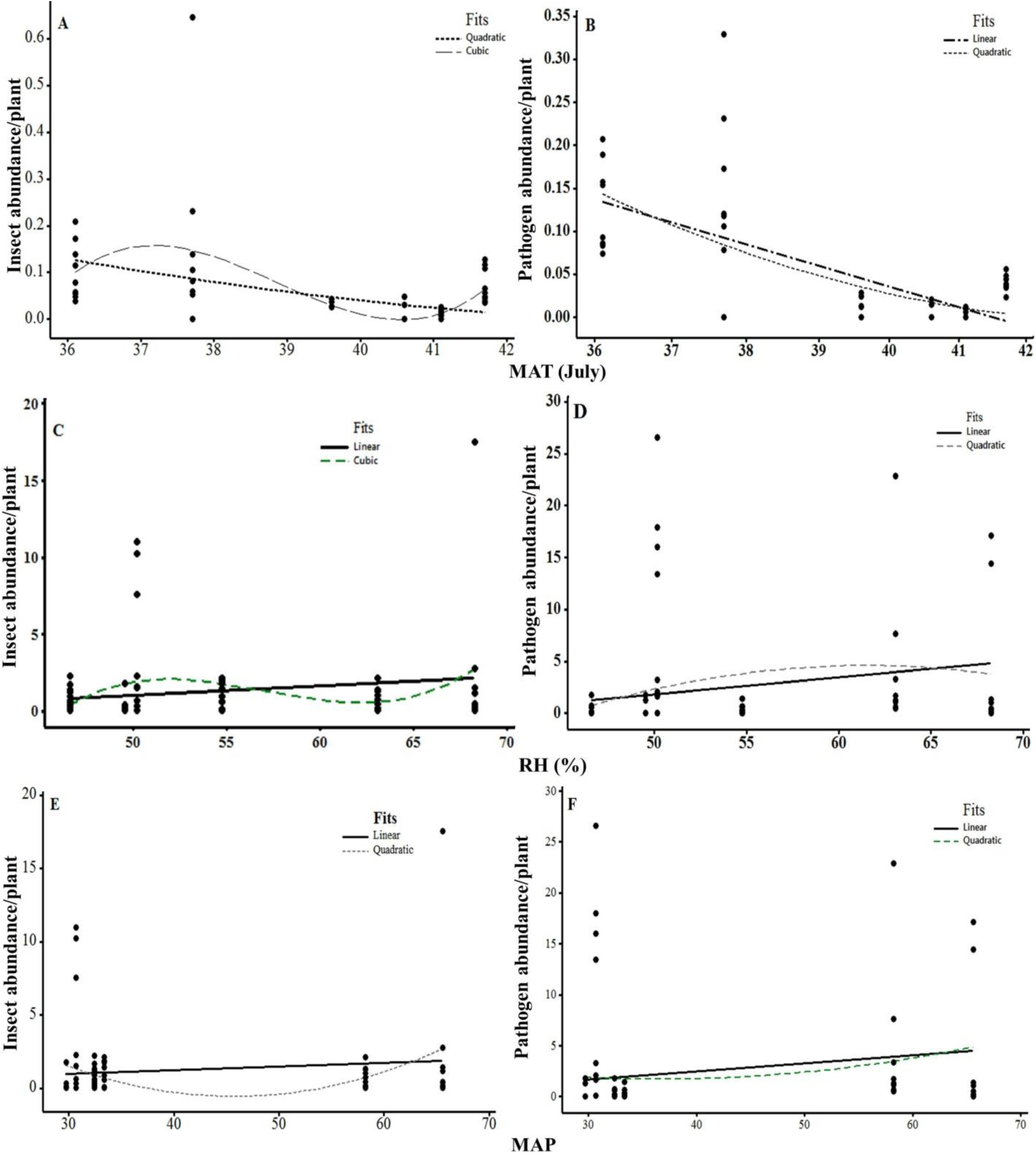
Scattered plot showing insect abundance per plant in a quadratic ring present on *Xanthium strumarium* with Maximum Annual Temperature (MAT) in July

The multiple comparisons of ecological factors demonstrated by MANOVA showed significant differences across the sites and invasion status of two natural enemies. The results of this experiment showed a significant (P<0.05) impact on the pathogen abundance, latitudinal, longitudinal gradients, MAP, MAT and RH (%) of the independent variables. Pathogen abundance did have a significant (P<0.05) positive effect on ecological factors. On the other hand, MANOVA regarding insect abundance differed significantly (P<0.05) with latitude and longitude and other environmental factors (MAP, MAT, and RH).

### 2.4. Model performance regarding principal component analysis

The second axis shows that the maximum insect and pathogen damage correlated negatively with the fourth axis. This third axis representing the trade-off between parasite, insect and pathogen abundance correlated positively between two components along with the fourth axis (Fig. 7A and 7C). In the PCA analysis, the first two axis explicated 71.39% and 20.96% of the total distinctions. On the other side, pathogen abundance, insect abundance, and parasite abundance correlated positively having 1, 3 and 4 axes. Insect damage showed a strong negative effect on the second component in PCA (Fig. 7B).

**Figure 7.**
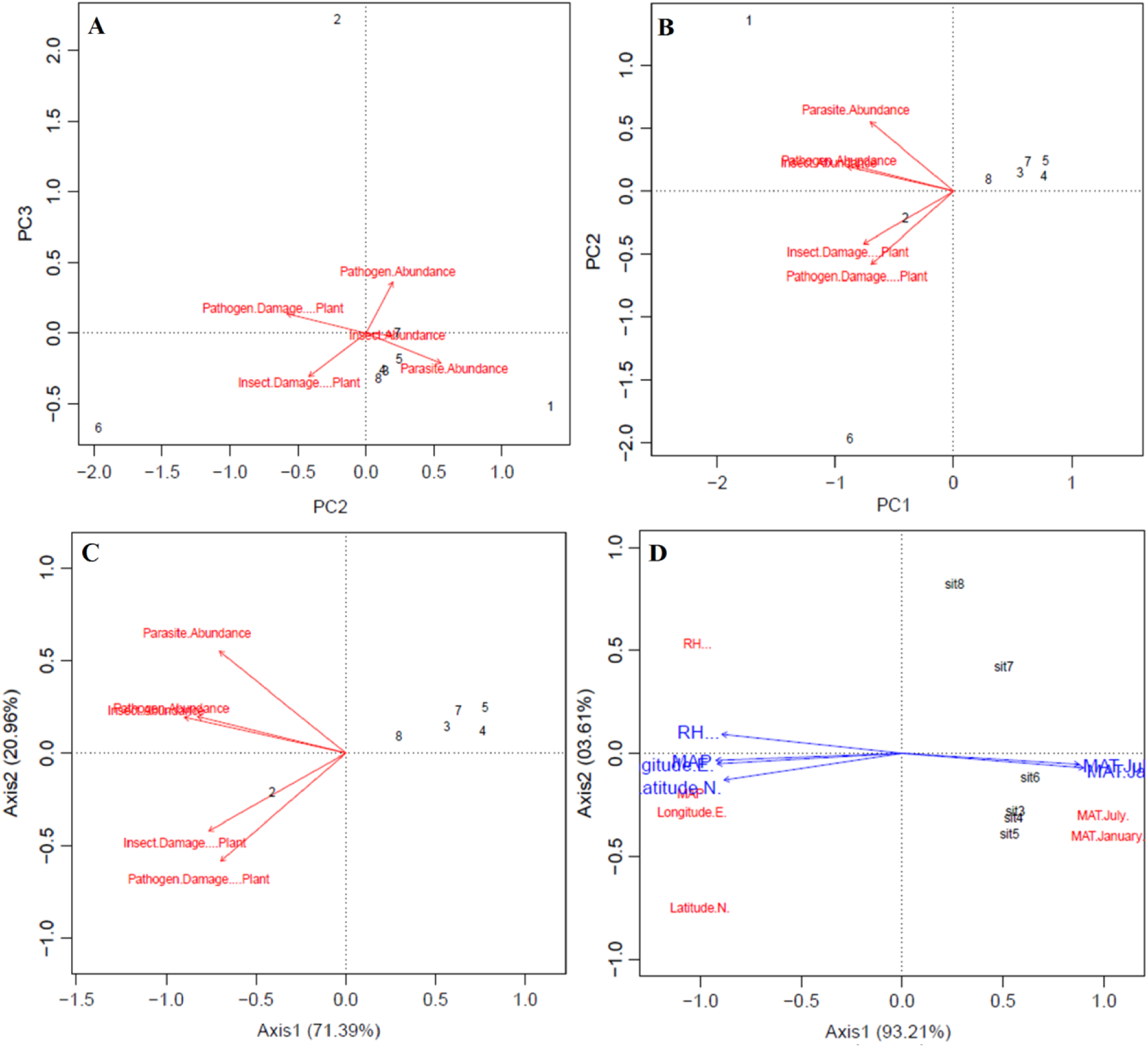
The plots showing the correlation between natural enemies abundance at different latitudes on *Xanthium strumarium* invasive weed.

In the PCA, the first two axis clarify 93.21% and 3.61% of the total deviation (Fig. 7D). The first axis showed mean maximum temperature in January and July correlated positively with the second axis representing the exchange among MAP, Latitude, longitude and correlated negatively compared to the first axis (Fig. 7). However, the third axis of RH (%) correlated positively with the site, which was present in fourth axis that also correlated positively or compared to other components. Therefore, the first component focuses on the positive correlation of temperature with natural enemy pressure on *Xanthium strumarium* in eight different sites in Northeast China.

### 2.5. Model performance regarding redundant analysis

In RDA one and RDA two correlations MAP, latitude and longitude were correlated negatively with natural enemy infestations. However, Site and MAT in July correlated positively in plant-herbivore interaction mechanism in axis 1. RH and site have a positive relationship in the third axis and the fourth axis. The RDA two and three correlation showed that relative humidity (%) was positively correlated with natural enemy infestations. MAP & MAT in July were correlated negatively with plant-herbivore interactions mechanisms along the second axis (Fig. 8A).

**Figure 8.**
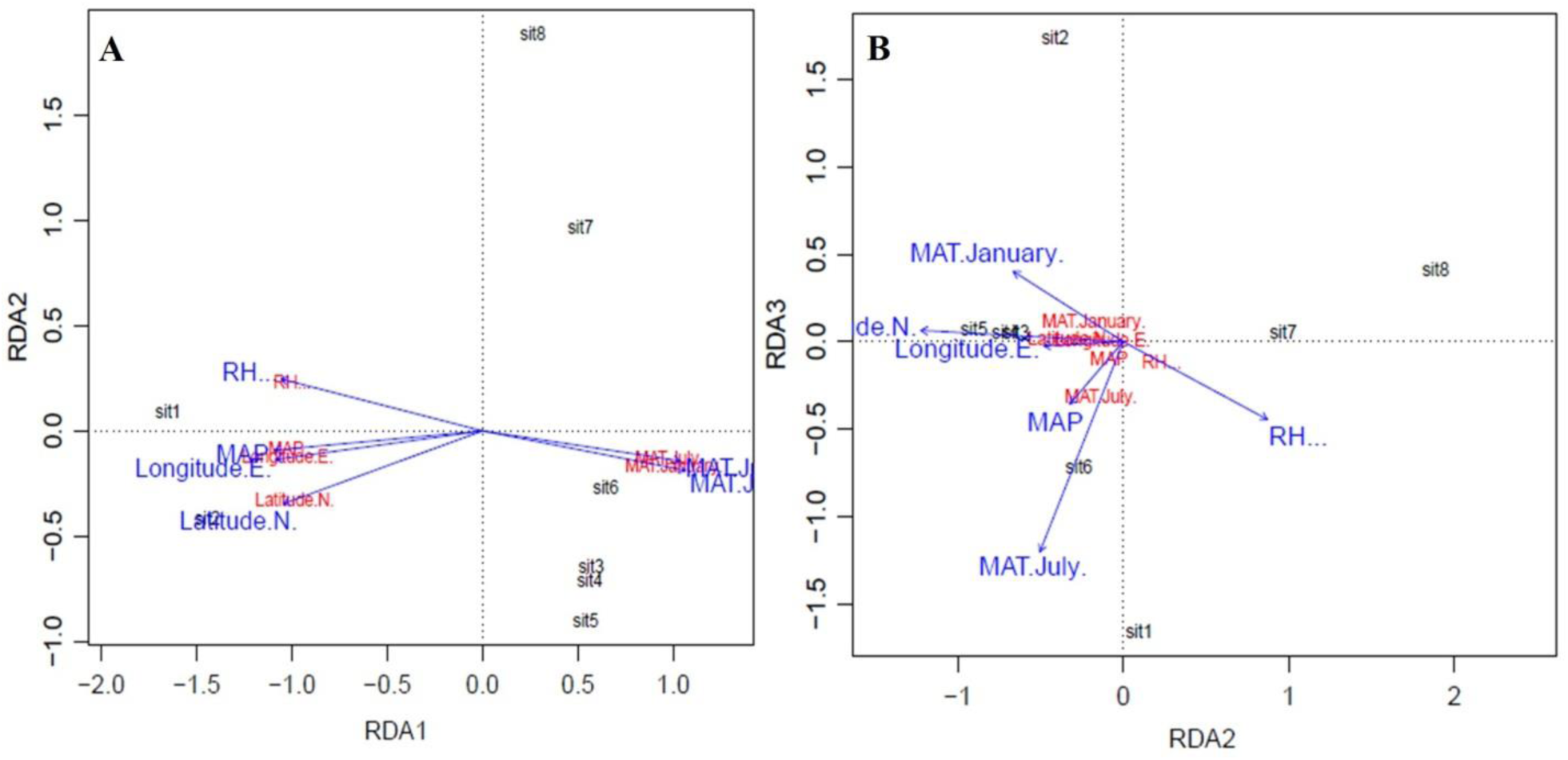
The correlation between the different abiotic factors, locations, as well as the first and second (**A** & **B**) component in RDA. The arrow shows dependent and independent variables present in each studied sites, RH means relative humidity (%); MAT means mean annual temperature, MAP means mean annual precipitation (mm) and it means different sites.

**Fig. 9.**
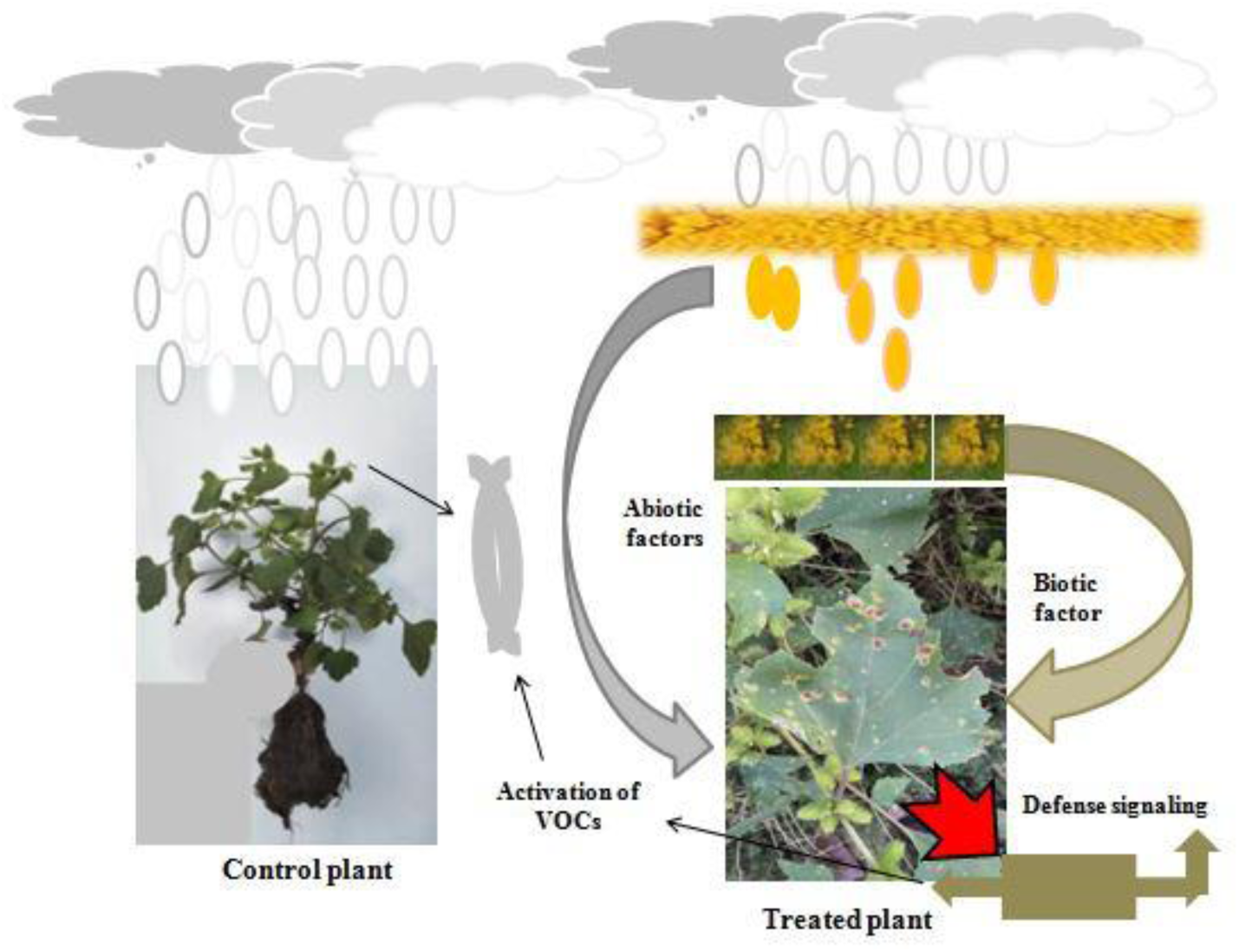
The presence of biological control agents in a natural ecosystems impact the performance of organisms that collectively involved in ecological niche interactions. VOC produce defensive signaling against pathogen attack in *Xanthium strumrium*

The plant height (cm) and diameter (mm) are correlated positively with the first axis in RD1 and RD3 analysis. The maximum value of the first and second axis described large variables that showed an abundance of natural enemy’s pressure in the ecosystem (Axis 2).

### 2.6. Solvents extraction and model fitness

The extraction was performed by high polarity (ethanol) and low polarity (distilled water) compounds from control and rust infected treatment. The first, second and third extractions were computed using each of 72 hours by maceration process. Extract yields (g) were calculated (Table 2) while the physical properties of these extracts were measured visually (Table 1). The results reported that maximum extract yield was recorded low polarity distilled water extracted treated area (0.191g) followed by control (0.155g) showed colored properties reddish brown, shiny, not sticky, crystalline and shiny, sticky and slippery respectively. Similarly the results suggested that extract yield was recorded by ethanol extracted produced (0.137g) in control followed by treated (0.137g) showed dark greenish with light greenish residues, shiny/less shiny, sticky, slowly flow able respectively (Table 2). These yields were statistically non significant (P > 0.05) to each other by using 5 gram original powdered leave sample.

**Table 1:** Co-efficient of determination model for cumulative extract yield (g) of invasive cocklebur.

| Sr. No. | Solvents extracts | *Xanthium strumarium* | Yield ± SD (gram) | Co-efficient of Determination | | Regression fit |
| --- | --- | --- | --- | --- | --- | --- |
| | | | | Regression fit | $R^2$ | RMSE |
| 1 | Ethanol | Control | 0.137±0.16a | -0.1483x + 0.4335 | 0.86 | 0.38 |
| 2 | | Treated | 0.079±0.06a | -0.0588x + 0.1971 | 0.99 | 0.19 |
| 3 | Distilled Water | Control | 0.155±0.09a | -0.085x + 0.3246 | 0.96 | 0.03 |
| 4 | | Treated | 0.191±0.15a | -0.1457x + 0.482 | 0.95 | 0.42 |

Whereas $R^2$ (Co-efficient of determination); RMSE (Root Mean Square Error); SD (Standard deviation); Control (Healthy leave); Treated (Rust Infected leaves); Same letter within a column indicate that mean values are differed non significantly ( $P > 0.05$ ) according to Tukey's HSD multiple range test

**Table 2:** Physical properties of Cockle bur (*Xanthium strumarium*) by different solvent extracts.

| Sr. No. | Solvents extracts | *Xanthium strumarium* | Physical properties of *Xanthium strumarium* leave |  |  |  |
| --- | --- | --- | --- | --- | --- | --- |
|  |  |  | Color | Opaqueness | Stickiness | Appearance |
| 1 | Ethanol | Control | Dark green/Greenish residues | Shiny | Sticky | Slowly flow able |
| 2 |  | Treated | Dark green/Light greenish residues | Less shiny | Sticky | Slowly flow able |
| 3 | Distilled Water | Control | Reddish brown/Light greenish residues | Shiny | Sticky | Slippery |
| 4 |  | Treated | Reddish brown/Reddish brown residues | Shiny | Non-sticky | Crystalline like brown sugar |

Whereas control (healthy leave); treated (rust infected leaves)

The coefficient of determination (*R*^*2*^) suggested that yield in response to *Xanthium strumarium* both in control and treated plants suggested maximum *R*^*2*^ (0.99), (0.96) respectively, however ethanol and distilled water recorded yield with RMSE (0.19) and (0.03) showed awesome model fitness.

### 2.7. Qualitative screening of phytochemicals

The qualitative screenings were used to identify bioactive constituents involved in pathogen defense mechanisms. The controlled and treated plants profiling confirmed the existence of essential ingredients such as flavonoids, alkaloids, terpens, phenols and saponins (Table 3-7) in these samples.

**Table 3:**
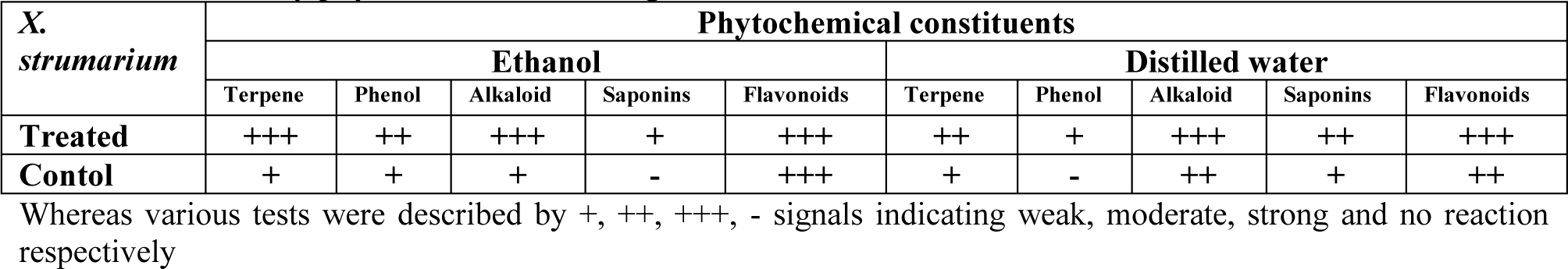
Preliminary phytochemical screening of leave extract of invasive *Xanthium strumarium*.

**Table 4:**
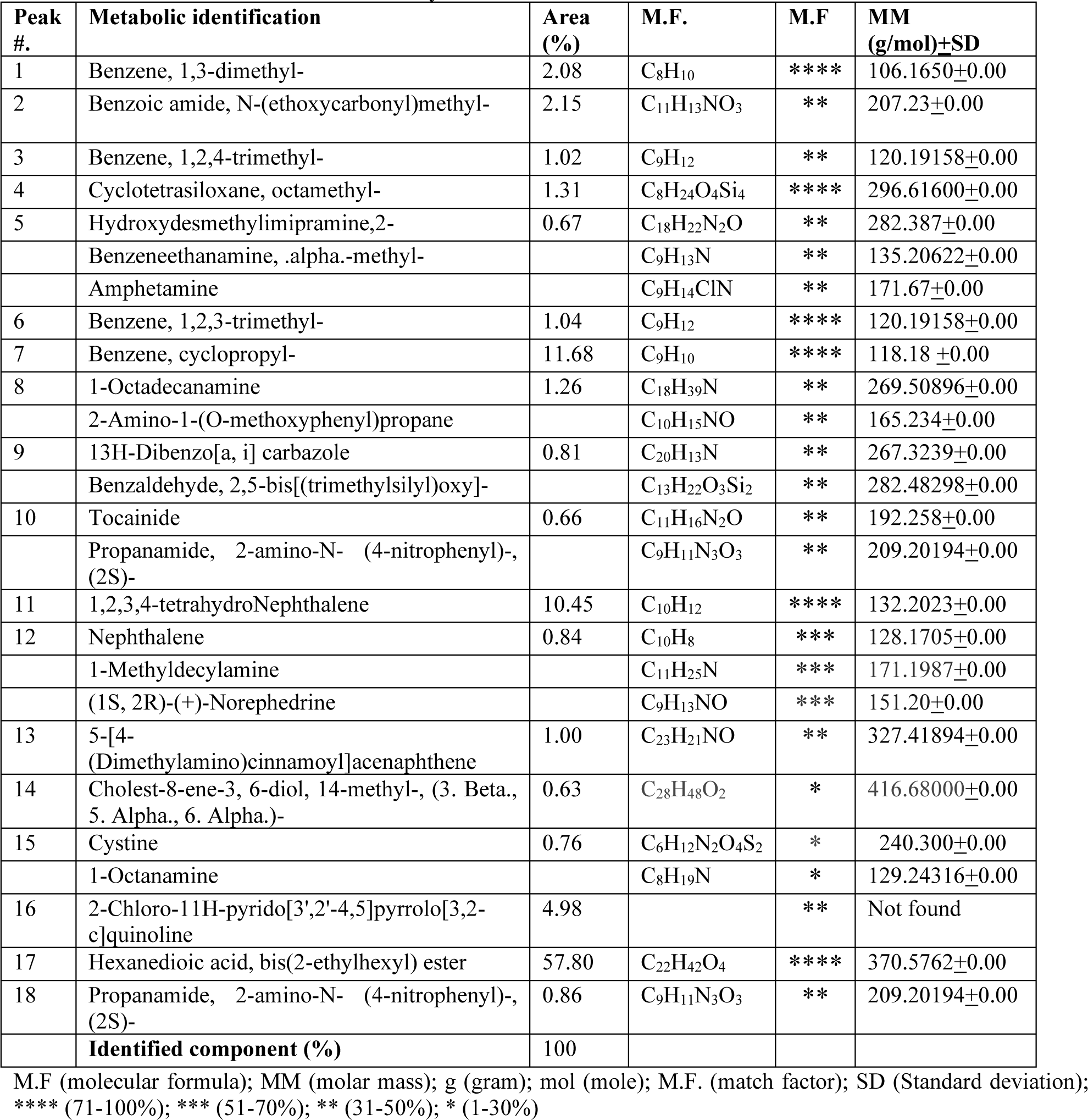
*Xanthium strumarium* extracted by Ethanol in control.

**Table 5:**
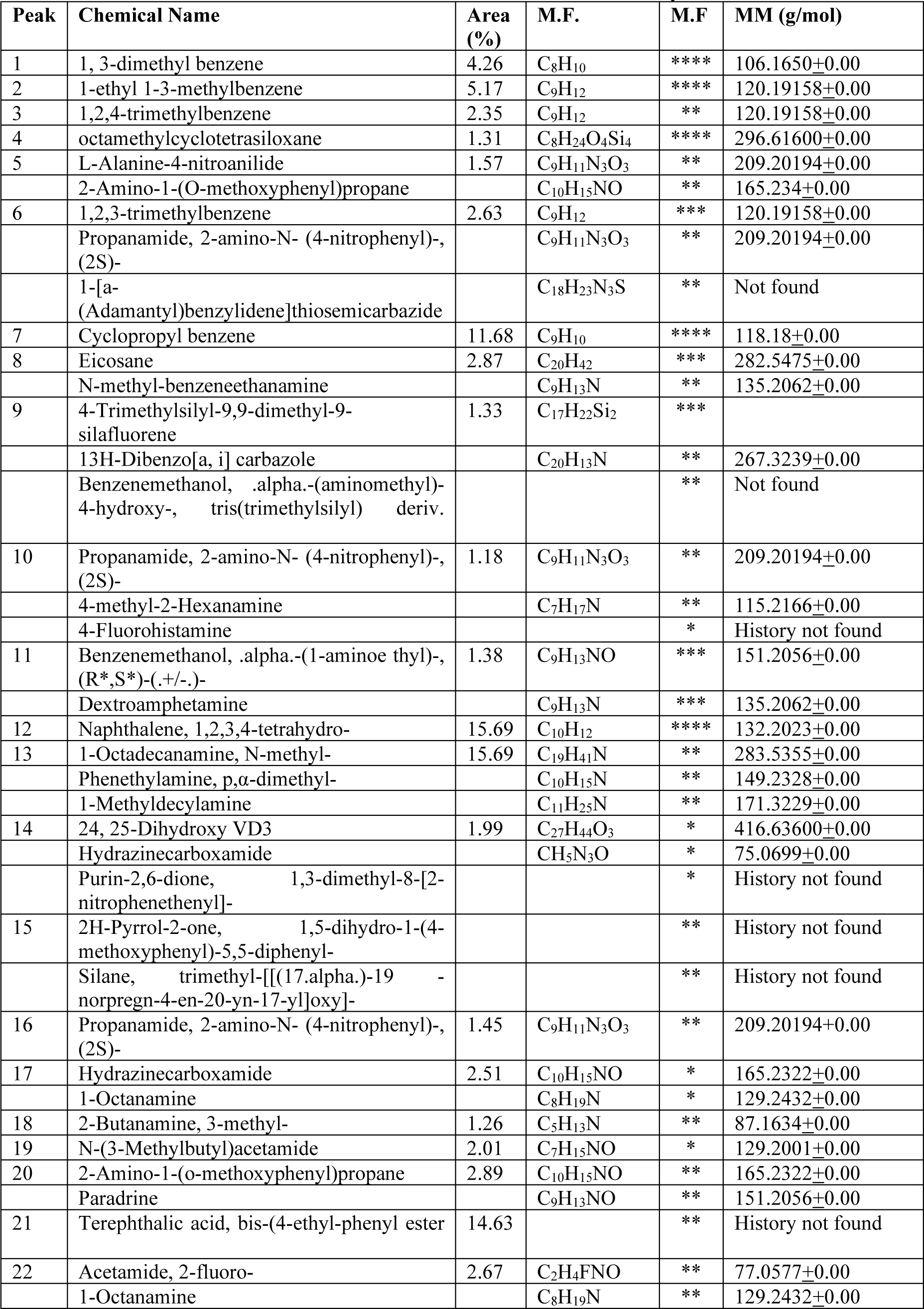
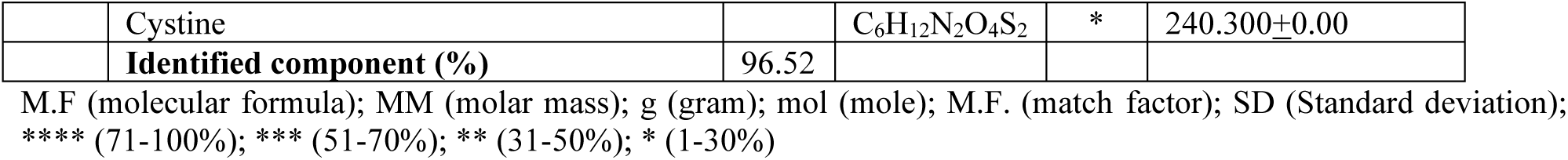
*Puccinia xanthii* infected *Xanthium strumarium* leaves extracted by Ethanol in treatment.

**Table 6:** *Xanthium strumarium* leaves extracted by Distilled Water (Control)

| Peak | Chemical Name | Area (%) | M.F. | M.F. | MM (g/mol) |
| --- | --- | --- | --- | --- | --- |
| 1 | Benzene, 1,3-dimethyl- | 5.34 | C<sub>8</sub>H<sub>10</sub> | **** | 106.1650±0.00 |
| 2 | 1-Methyldecylamine | 1.26 | C<sub>11</sub>H<sub>25</sub>N | ** | 171.3229±0.00 |
|  | 2-Aminononadecane |  | C<sub>19</sub>H<sub>41</sub>N | ** | 283.5355±0.00 |
| 3 | Benzene, 1-ethyl-3-methyl- | 5.03 | C<sub>9</sub>H<sub>12</sub> | **** | 120.1916±0.00 |
| 4 | Benzene, 1,2,3-trimethyl- | 2.10 | C<sub>9</sub>H<sub>12</sub> | ** | 120.1916±0.00 |
| 5 | Cyclotetrasiloxane, octamethyl- | 2.73 | C<sub>8</sub>H<sub>24</sub>O<sub>4</sub>Si<sub>4</sub> | **** | 296.6158±0.00 |
| 6 | 2-Butanamine | 1.46 | C<sub>4</sub>H<sub>11</sub>N | ** | 73.1368±0.00 |
|  | 1-Octadecanamine, N-methyl- |  | C<sub>19</sub>H<sub>41</sub>N | ** | 283.5355±0.00 |
| 7 | Benzene, 1,2,4-trimethyl- | 2.96 | C<sub>9</sub>H<sub>12</sub> | **** | 120.1916±0.00 |
| 8 | 2-Butanamine | 1.27 | C<sub>4</sub>H<sub>11</sub>N | ** | 73.1368±0.00 |
|  | 2-Amino-1-(O-methoxyphenyl)propane |  | C<sub>10</sub>H<sub>15</sub>NO | ** | 165.2322±0.00 |
|  | Phenylpropanolamine |  | C<sub>9</sub>H<sub>13</sub>NO | ** | 151.2056±0.00 |
| 9 | Indane | 28.61 | C<sub>9</sub>H<sub>10</sub> | **** | 118.1757±0.00 |
| 10 | 1-Octadecanamine | 3.43 | C<sub>18</sub>H<sub>39</sub>N | ** | 269.5090±0.00 |
|  | Benzeneethanamine, N-methyl- |  | C<sub>9</sub>H<sub>13</sub>N | ** | 135.2062±0.00 |
| 11 | Benzaldehyde, 2,4-bis(trimethylsiloxy)- | 1.43 |  | *** | not Found |
|  | 1,5-Diphenyl-3-methylthio-1,2,4-triazole |  |  | ** | not Found |
| 12 | 3,4-Methylenedioxy-amphetamine | 2.00 | C<sub>10</sub>H<sub>13</sub>NO<sub>2</sub> | ** | 179.2157±0.00 |
|  | Phenol, 4-(2-aminopropyl)-, (.+/-.)- |  | C<sub>9</sub>H<sub>13</sub>NO | ** | 151.2056±0.00 |
| 13 | Benzeneethanamine, N-methyl- | 1.71 | C<sub>9</sub>H<sub>13</sub>N | ** | 135.2062±0.00 |
|  | 2-Amino-1-(O-methoxyphenyl)propane |  | C<sub>10</sub>H<sub>15</sub>NO | ** | 165.2322±0.00 |
| 14 | Naphthalene, 1,2,3,4-tetrahydro- | 16.48 | C<sub>10</sub>H<sub>12</sub> | **** | 132.2023±0.00 |
| 15 | Silane, trimethyl-[[[(17.alpha.)--norpregn-4-en-20-yn-17-yl]oxy]- | 3.17 |  | ** | not Found |
|  | 5-[4-(Dimethylamino)cinnamoyl]acenaphthene |  | C<sub>12</sub>H<sub>8</sub> | ** | 152.196±0.00 |
|  | Dibenz[b,d]cycloheptane, 2,4,7-trimethoxy-11-acetamino- |  | C<sub>21</sub>H<sub>25</sub>NO<sub>5</sub> | ** | not Found |
| 16 | Phenethylamine, p,.alpha.-dimethyl | 1.31 | C<sub>10</sub>H<sub>15</sub>N | ** | 149.2328±0.00 |
|  | n-Hexylmethylamine |  | C<sub>7</sub>H<sub>17</sub>N | ** | 115.2166±0.00 |
|  | 2-Octanamine |  | C<sub>8</sub>H<sub>19</sub>N | ** | 129.2432±0.00 |
| 17 | Rhenium, pentacarbonyl(trifluorovinyl)- | 1.61 |  | * | not Found |
|  | Phosphine oxide, bis(pentamethylphenyl)- |  |  | * | not Found |
|  | 4-Hexenethioic acid, 3-(t-butyl dimethylsilyloxy)-, S-t-butyl ester |  |  | * | not Found |
| 18 | Benzeneethanamine, N-methyl- | 1.29 | C<sub>9</sub>H<sub>13</sub>N | ** | 135.2062±0.00 |
|  | 1-Octadecanamine, N-methyl- |  | C<sub>19</sub>H<sub>41</sub>N | ** | 283.5355±0.00 |
| 19 | 1-Octanamine | 1.69 | C<sub>8</sub>H<sub>19</sub>N | * | 129.2432±0.00 |
|  | Acetamide, 2-fluoro- |  | C<sub>2</sub>H<sub>4</sub>FNO | * | 77.0577±0.00 |
| 20 | 1-Octadecanamine, N-methyl- | 2.57 | C<sub>19</sub>H<sub>41</sub>N | ** | 283.5355±0.00 |
|  | 2-Amino-1-(O-methoxyphenyl)propane |  | C<sub>10</sub>H<sub>15</sub>NO | ** | 165.2322±0.00 |
|  | 2-Hexanamine, 4-methyl- |  | C<sub>7</sub>H<sub>17</sub>N | ** | 115.2166±0.00 |
| 21 | Phenol, 4-(3,4-dihydro-2,2,4-trimethyl- | 10.20 | C<sub>18</sub>H<sub>20</sub>O<sub>2</sub> | ** | 268.3502±0.00 |
|  | 2H-1-benzopyran-4-yl)- |  |  |  |  |
| 22 | 1-Octadecanamine, N-methyl- | 2.33 | C<sub>19</sub>H<sub>41</sub>N | ** | 283.5355+0.00 |
|  | **Identified component (%)** | 99.98 |  |  |  |

M.F (molecular formula); MM (molar mass); g (gram); mol (mole); M.F. (match factor); SD (Standard deviation); \*\*\*\* (71-100%); \*\*\* (51-70%); \*\* (31-50%); \* (1-30%)

**Table 7:**
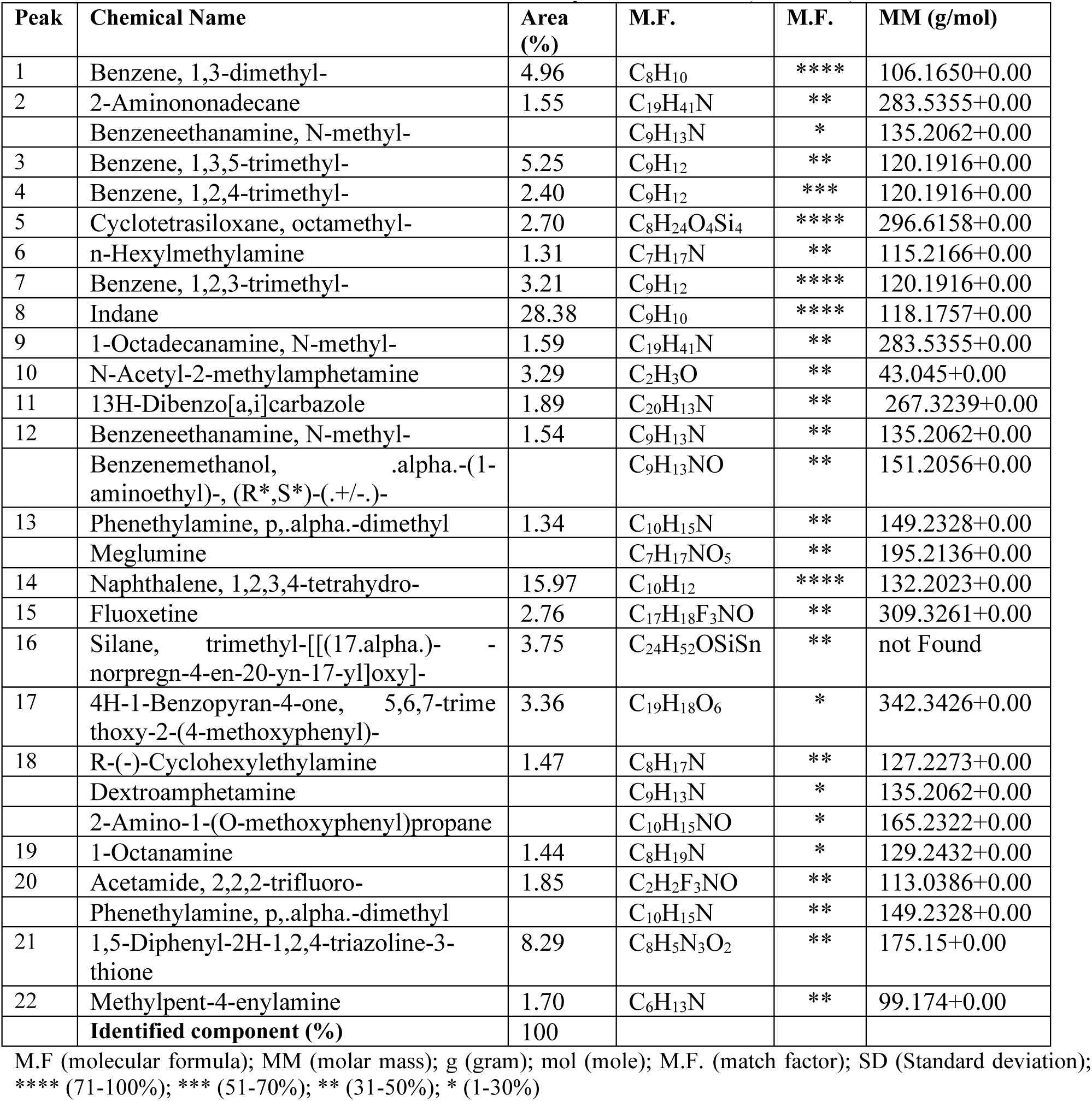
*Xanthium strumarium* infected leaves extracted by Distilled Water (treatment)

### 2.8. Chemical compositions of invasive cocklebur

The chemical compositions of the *Xanthium strumarium* controlled leave (healthy) extraction was performed by ethanol determined by GC-MS analytical technique with area (%), molecular formula (M.F) and molecular weight (M.W) were determined. Eighteen with eight sub-compounds (Table 3) were identified having monoterpenes; aldehydes; diterpenes and esters; however, the following compounds are representing the major constituents: (17) Hexanedioic acid, bis(2-ethylhexyl) ester (57.80%), (7) Benzene, cyclopropyl- (11.68%), (11) 1,2,3,4-tetrahydro Nephthalene (10.45%), (16) 2-Chloro-11H-pyrido[3’,2’-4,5]pyrrolo[3,2-c]quinoline (4.98%), (2) Benzoic amide, N-(ethoxycarbonyl)methyl- (2.15%), (1) Benzene, 1,3-dimethyl-(2.08%) respectively having diversified match factors. On the other hand pathogen *Xanthium strumarium* recorded twenty two and fifteen sub-compounds (Table 4) were identified having monoterpenes; diterpenes and polyterpene however, the following compounds are representing the major constituents: (12) Naphthalene, 1,2,3,4-tetrahydro- (15.69%), (13) 1-Octadecanamine, N-methyl-; Phenethylamine, p,α-dimethyl-; 1-Methyldecylamine (15.69%), (21) Terephthalic acid, bis-(4-ethyl-phenyl ester (14.63%), (7) Cyclopropyl benzene (11.68%), (2) 1-ethyl 1-3-methylbenzene (5.17%), (1) 1, 3-dimethyl benzene (4.26%) respectively having diversified match factors. The low polarity solvent extraction and chemical compositions of the *Xanthium strumarium* controlled leave recorded twenty two and eighteen sub-compounds (Table 5) were identified having monoterpenes; phenols, esters, and other compounds however, the following compounds are representing the major constituents: (9) Indane (28.61%), (14) Naphthalene, 1,2,3,4-tetrahydro- (16.48%), (21) Phenol, 4-(3,4-dihydro-2,2,4-trime thyl-2H-1-benzopyran-4-yl)- (10.20%), (1) Benzene, 1,3-dimethyl- (5.34%), (3) Benzene, 1-ethyl-3-methyl- (5.03%), (10) 1-Octadecanamine (3.43%) respectively having diversified match factors. The chemical compositions of the *Xanthium strumarium* infected leave recorded twenty two and seven sub-compounds (Table 6) were identified having monoterpenes; deterpenes; phenols, esters, and other compounds however, the following compounds are representing the major constituents: (8) Indane (28.38%), (14) Naphthalene, 1,2,3,4-tetrahydro- (15.97%), (21) 1,5-Diphenyl-2H-1,2,4-triazoline-3-thione (8.29%), (3) Benzene, 1,3,5-trimethyl- (5.25%), (1) Benzene, 1,3-dimethyl- (4.96%), (16) Silane, trimethyl-[[(17.alpha.)-norpregn-4-en-20-yn-17-yl]oxy]- (3.75%), (17) 4H-1-Benzopyran-4-one, 5,6,7-trime thoxy-2-(4-methoxyphenyl)- (3.36%), (10) N-Acetyl-2-methylamphetamine (3.29%), (7) Benzene, 1,2,3-trimethyl- (3.21%) respectively having diversified match factors while other compounds were present in the minor quantities.

## 4. Discussion

During an extensive field survey at eight locations in Liaoning and Hebei Province, it was found that in *X. strumarium* herbivore interactions some growth traits were correlated with latitudinal gradients. It is stated that plant-herbivore interactions happen in every ecosystem that afforded a key opportunity for energy stream to high trophic levels. This is an ancient theory to clarify the different levels of latitudes in plant range suggested comparatively constant and warm climate led to strong biotic interactions in contrast to temperate regions. The natural enemy pressures have been evaluated across different levels of latitudes to trial out biotic interactions hypothesis [50]. Latitudinal trends involve a deviation in abiotic (daylight period, temperature fluctuations) and biotic (plant species diversity) conditions and contain significant effects in several plant traits (leaf quality, productivity).

### 4.1. Factor affecting the growth of *X. strumarium*

According to latitudinal trends for the size of the plant response, the smaller size was observed at elevated latitudes than inferior latitudes under a low positive growing situations [51]. This type of latitudinal pattern in term of plant growth characteristics was described in toxic plants [52,53]. The results of this experiment was in accordance to the researchers who reported that plant-insect-pathogen (PIP) interactions influence on the plant performance and growth related factors [54]. It was also investigated that PIP played a significant impact to reduce height, branching and biomass [55,56] of the plants. Contrarily some studies found an insignificant difference (P>0.05) in latitudinal trends regarding plant size in *Phytophthora americana* [57]. The current study indicated that plant height (cm) increased significantly at 41°N than at 40°N latitude. One clarification for latitudinal mold is the maximum rainfall at 41°N compared to 40°N resulting in enhanced plant height (cm) of *X. strumarium* species.

There were some positive changes in *X. strumarium* plant height (cm) with increasing latitudinal patterns. Plant growth parameters correlated positively and increased with the elevated trend of latitudes from 40°N to 41°N. This latitudinal pattern supports the study by Stutzel [58]. Moreover, it was found that *Phytolacca americana* showed a small stem diameter (mm) at higher latitudinal gradients, suggesting style adaptation approach [57]. The findings of the present study suggested that the stem diameter of *X. strumarium* reduced significantly with increased longitudes. This situation might be possible to the maximum insect abundance in relation to damage at this latitude. Future investigations are recommended to deeply incorporate environmental, latitudinal, longitudinal factors to filter out the influence of abiotic dynamics on toxic weeds over expanded geographical areas to afford a significant prediction for wide distribution. The present results are in accordance with studies that reported that growth parameters are significantly related to latitudinal gradients.

In this study the height (cm) of *X. strumarium* was increased significantly at 41°N and the results were in accordance to the researchers who reported that the stem diameter of the toxic weed increased at north latitudinal patterns (42°N to 32°N) which was consistent to our findings that related to natural enemy abundance at northern populations [59]. These results are in accordance to the scientists who reported that slight variation in temperature resulting in a maximum difference in biological control agents than latitudinal pattern, however, in this situation rainfall and population density were less important than other ecological factors [60].

### 4.2. Response of *E. strenuana* and *P. xanthii* abundance with latitudes

The finding indicated that natural biological control agents were generally ten times more abundant at 31-36°N than 41-44°N [4]. The experiment described that *Epiblema* was recorded a maximum of 7.3 times more abundant at 41.51279°N. The results investigated that natural enemy pattern was correlated with a latitudinal gradient. These results were in accordance with studies reported that *Lipara rufitarsis* (gall fly) on native *Phragmites australis* (wetland grass) lineage increased from 27% at 36.5°N to 37% in northern populations (43.6°N) [4]. But, it was suggested that latitudinal patterns in plant natural enemy interactions are a universal event across a variety of invaded classifications [61]. It was also suggested that a significant latitudinal pattern described a less damage rate at low latitude [14], which is in line with the results of the present investigations.

The results do not support the studies found that the pressure of biological control agents in logically happening sites of *P. australis* is maximum at low latitudes [17]. The present results strongly support the hypothesis that natural enemy abundance can be a selective vigor contributing to the experiential mold of increased and decreased plant palatability at low versus high latitudes [5]. The relationship of insect abundance with latitudes was in line with the previously reported studies that latitudinal gradients associated with plant natural enemy relations are phenotypically plastic [18]. However, invasive are more plastic than non invasive weeds [18,62,63]. The pattern of latitudes might be dependent upon biotic and abiotic relationships [61]. The results of this experiment reject the claim that the stem borer damage increases with decreasing latitudes [60]. The experiment studied deviation in abundance and relationship with leaf-chewing herbivores on *Spartina alterniflora* suggested that the value of elevated latitudes having *S. alterniflora* plants are extra palatable to natural enemies compared to low latitudes [64]. The present experiment suggests that *X. strumarium* at high latitude, 41.51279°N area is more palatable to *Epiblema strenuana* compared to low latitude areas.

### 4.3. Response of *E. strenuana* and *P. xanthii* with abiotic factors

Previously, it was believed that natural enemy pressure decreases significantly with latitude. Now, experimental investigations have unproductive to advocate this hypothesis leading to obsessive debate. In accumulation and providing a quantitative measurement, the researchers tested the latitude-herbivory theory and the connection among temperature and rainfall to herbivory [65]. Based on the literature survey, the results suggested that natural enemy abundance reduced with latitudes and elevated with the temperature in the North regions. In the experiment on the northeastern area of China, this hypo theoretical study is reciprocal to the current study. The role of environmental factors with latitudes in plant natural enemy interactions has not been evaluated [8,50]. The researchers investigated temperature (°C) and rainfall (mm) are inter connected with latitude that play a significant role in plant natural enemy interactions mechanism [66]. Recent investigations confirmed the correlation of leaf traits with environmental factors [67]. Temperature can affect the visible morphological characters, survival, abundance, and distribution of the biological control agents [66]. Rainfall, annual temperature range were the strong forecaster linked with *P. solenopsis* diversification, however, precipitation of cold part was negatively associated with *P. solenopsis* incidence [68]. Herbivory and pathogen have contrasting, independent effects on plant growth, with pathogen decreasing and herbivory increasing growth characteristics of the plant [69]. Nevertheless, in this experiment, *Epiblema* did not show its relationship with precipitations, but it was revealed that *P. xanthii* depends upon precipitation and relative humidity for its spreading. On the other hand, the insects oviposit at night are principally dependent on nighttime temperatures for oviposition. For foliar fungi, infection and sporulation frequently involve close to 100% RH, and these circumstances happen mainly during overnight dewfall [70]. Therefore, an optimum temperature for their biological activity in this specific time is principally significant [71], however, the temperature has been pointed out a significant driver of plant natural enemy inter relationship [66,72,73].

The studies reported that plant natural enemy interaction with latitudinal pattern recorded higher herbivore abundance at lower latitudes. The negative relationship between herbivore abundance and latitude were found in the present study, which supports the findings of other studies [4,10,74]. In addition, temperature thresholds could reduce abundance at higher latitudes in subtropical areas [75]. It is a phenomenon that insect infestation is maximum at a warmer climate than colder climatic conditions [14,76]. However, the results of the present study became contrary to this phenomenon, which suggested that damage varied with insect species and its distributions at different latitudes. The abundance of *Epiblema* and *P. xanthii* was maximum at 36 °C (41°N) compared to warmer temperatures regions up to 41 °C (40°N). Moreover, the researchers incorporated fluctuations in temperature and precipitation rather than plant natural enemy interactions with latitude which plays a significant role in understanding the impacts of abiotic and biotic interactions [7].

### 4.4. Qualitative screening of phytochemicals

Initial phytochemical screening investigated the presence of flavonoids, phenols, saponins, alkaloids, terpens, silicon and sulpher containing compound (Table 3).

### 4.5. Abundance in defensive behavior

The plant did not escape from biological control agents (BCA’s) already present in natural ecosystems. However, after herbivore attack the plants produced VOC’s used as defense mechanisms. These are sub-divided into three major groups of chemicals known as terpens, phenols and N compounds. The effects of terpenes are toxic played a negative impact on herbivores. The lignin and tannins are phenols present in cell wall, however, the alkaloids having N present in larger area in plants. In rangeland ecosystem, invasive weeds are present and may consume by natural animals caused failure in nervous system resulting paralysis and even sudden deaths was investigated [77].

### 4.6. GC-MS profiling of *X. strumarium* leave

Under favorable environmental conditions, plants are constantly exposed to a wide range of biotic interactions (pathogens). The present study revolved the GC-MS of leave described the metabolic identification of high polar extraction solvent (ethanol) suggested that monoterpene covered an area (11.29%) in control treatment with medium similarity compared to treated (rust infected) produced monoterpene (15.69%) with significantly high match factor. Similarly low polar solvent (distilled water) described monoterpene (10.48%) in control compared to pathogen infected (15.97%) Xanthium strumarium infected leaves. These results suggested that the plant secreted chemicals in the form of carbon-hydrogen (CH) compounds which were derived from terpenoid and ethylene produce significant abundance (%) compared to control played a significant role in plant-herbivore defense mechanism. These results are in line with the researchers who reported that rate of terpene emission and composition against the natural herbivore enhanced modes of defense [78] in the weed plants. In the present study polyterpene is present in the pathogen treated GC-MS results compared to the control treatment. These results are in line with the researchers who reported that polyterpene provides protection mechanism and healing affect as a defense against herbivores [79,80].

#### 4.6.1 Flavonoids

Flavonoids are the largest classes of phenols, present in the epidermal cells of the plants [81]. These compounds are played important role defense mechanism against pathogens and included like flavenol, flavone, flavanone, flavan-3-ol, isoflavone, and anthocyanin.

#### 4.6.2. Phenol compounds

The results of GC-MS suggested that *Puccinia xanthii* infected (treatment) leaves covered maximum area (%) compared to control treatments (Table, 3-7) due to breakdown of the chemical compounds that proved the hypotheses that volatile organic compounds altered infrastructures of the leave chemistry that led to activeness of plant defensive chemicals. These results are in accordance to the scientists who reported that phenolic compounds involved in resistance mechanism against pathogens and other herbivores. Several classes of phenolics have been categorized on the basis of their basic skeleton: C6-C1 (phenolic acid), C6-C2 (acetophenone, phenylacetic acid), C6-C3 (hydroxycinnamic acids, coumarins, phenylpropanes, chromones), C6-C4 (naphthoquinones), C6-C3-C6 (flavonoids, isoflavonoids), (C6-C3-C6)2 (biflavonoids), (C6-C3)n (lignins), (C6-C3-C6)n (condensed tannins). Some phenolic compounds are responsible for the inhibition of fungal growth. Chemical ecology of plants described that phenols are capable in resistance mechanisms against fungal pathogens [82-87], however phenylalanine ammonia lyase activity was increased in wounded plant tissues [88].

#### 4.6.3. Nitrogen Secondary Products

In these products nitrogen is present acts as defensive elements against herbivores [77]. This group include alkaloids which is a very large chemical group that is found in about 20% of vascular plant species [89]. The plant defense responses against pathogens are determined by the time, nature of infection [90] however, these responses are typical [91,92] and phenotypically plastic that can be altered due to biotic stress. The evidence advocated that the involvement of jasmonic acid (JA), ethylene (ET), SA and abscisic acid (ABA) are also VOC’s used in plant defense [42,93,94]. The pathogen damage symptoms identified in *X. strumarium* leave tissues that stimulate the emission of VOC’s. However in our experiment high pathogen pressure (8.97%) investigated on the leaves tissues gave the evidence and some accumulated results in the last few years indicating that volatiles produced from vegetative plant parts can be involved in plant fitness. Therefore in order to determine the plant-pathogen-defense mechanism, multi spectral research is needed on molecular and genetic level [95].

## 5. Future prospects

This study provides an evidence-based indication that natural enemies’ pressure varies with latitude in different areas. In the current experiment, *Epiblema strenuana* and *Puccinia xanthii* are the natural enemies having biological control potential against *Xanthium strumarium* recorded significant abundance and damage (%) at 41.51279°N latitude at 36°C temperature compared to 40.124301°N. The consideration of defense mechanism in plant is a key challenge because pathogen infections involved in profound physiology, morphology, anatomy and chemical variation processes in interaction mechanisms. Therefore, the ecologists, plant pathologists are advised to study this phenomenon in a more holistic way, integrating genetics, ecology and physiology to depict these complex interactions. The researchers are advised to explore the defensive mechanism of the plant to study the interaction of Jasmonic acid with insects, their pathway analysis. The significance of non-parallel different levels of latitudes for toxic weeds remain unexplored for other species interactions, mutualism, competitions which may play a vital role in amplification of invasion success in future.

## 6. Materials and Methods

### 6.1 Study locations

The study vicinities are the northern part of China, including the hilly area that edges South Korea. In these areas, the Shenyang Zone extends from 41.49490°N latitude and 123.35397°E longitudes. The maximum temperature ranges between 8.30 and 36.10°C and the minimum ranges between −32.90 and 12.40°C, but the annual rainfall was 698.5 mm. Due to the dynamic climate and the combination of hilly areas, these regions are rich biodiversity of biological control agents (BCA’s).

### 6.2. Herbivores, environment data collections

The ecological survey was conducted in eight different sites of Liaoning and Hebei Province (Fig. 1) by the line transects ecological method at field conditions during 2018. *X. strumarium* was attacked naturally by Rust (*P. xanthii*) and *E. struneuna* infestations were identified from the leaves. The transect line was sixty meters long parallel to the road or riverside, this line was divided into three similar portions each of twenty meter long collected randomly and data were recorded using quadratic ring [96-98] ring (100×100 cm) according to standardized protocol [99]. Three whole cocklebur plants in each quadratic ring were selected randomly for measuring their growth parameters and other characteristics. From each plant chosen in a quadratic ring, nine leaves were taken randomly from upper, middle and lower portions to calculate the damage caused by insect and pathogen and this procedure was repeated three times in a site. The latitude and longitude, minimum and maximum temperature, precipitation, and relative humidity (%) were recorded from the Chinese database. Location and elevation data were extracted from the digital elevation model [100]. The symptoms of damage caused by larva on leaves and *Puccinia xanthii* (Fig. 2) were identified by telia and teliospores [101,102]. The insect attack (%) was measured directly on the leaf tissues by ImageJ. However, pathogen damage (%) was measured first by adobe photoshop and then by using digital image processing approach [103,104].

The damage abundance caused by *Epiblema strenuana* and *Puccinia xanthii* on *Xanthium* leave, Simpson’s Index of Dominance (SID), Prominence Value (PV) in an ecosystem were calculated [105-109].

### 6.3. Invasive cocklebur collection

#### 6.3.1. Material preparations

The study was conducted in Shenyang and the leaves samples of *Xanthium* was collected from different areas in Shenyang and washed it gently first with tap water then in distilled water. The leaves were dried in the room temperature until all the water molecules were evaporated and the leaves became well dried keeping in view the grinding conditions. The leaves were grinded firstly in the local grinder, after that the material was shifted to the GT-200 grinder (Company) keeping the speed 1500rpm for 5 minutes (Figure 10).

**Figure 10:**
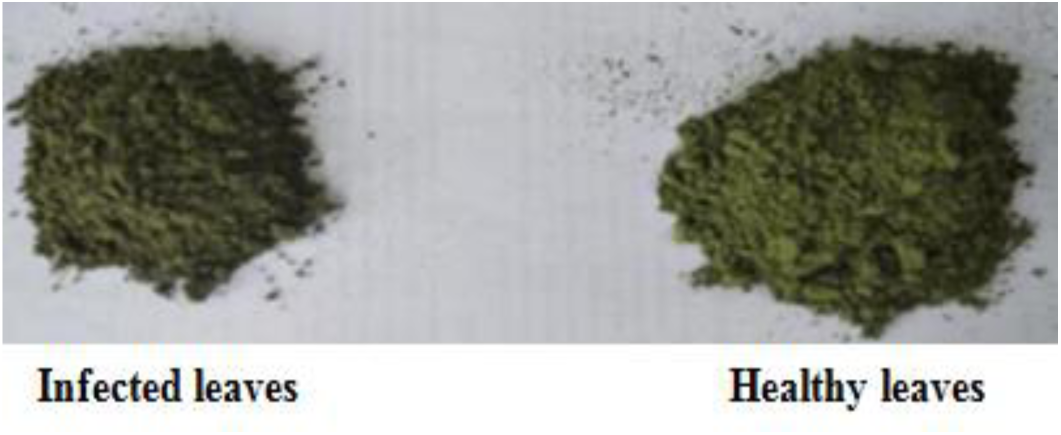
Cocklebur grinded healthy (control) and infected leaves (treated)

#### 6.3.2. Solvent extraction

According to polarity basis the powdered plant leaves were used for organic fractions [110]. Ethanol and distilled water @ 4ml g^−1^ of sample was used at room temperature for 72 hours [111-113]. Extracts were filtered and concentrated through the process of evaporation at room temperature. After complete evaporation the sample was weighed and first yield was calculated; however the same procedure was adopted for thrice to make sure that the whole polar and non-polar materials were eluted finally [114] stored at 4°C in air tight sterile tubes [115]. Physical properties of each extract (color, stickiness and appearance) recorded visually (Table 1) [116].

#### 6.3.3. Phytochemical screening

Phytochemical analysis were assessed according to the standards protocol to find out presence of various chemical compounds such as alkaloids, saponins, terpenoids and steroids [117], glycosides, flavonoids and flavones [118,119] phenols and tannins using different solvents [115,116,120]

### 6.4. Model validations

The relationship between treatments was investigated by co-efficient of determination (*R*^*2*^) values with root mean square error (RMSE). The values of *R*^*2*^ and *RMSE* represent the standard deviation of the prediction errors. Residuals are the measures that how the values are far away from the regression lines around our data points [121], that measures the best fitness of model validity [122,123]. In this experiment RMSE gave yield performance regarding range of model fitness [115,124].

### 6.5. Sample Preparation

Approximately 10mg of dry sample of each solvent extract was accurately weighed taken in the centrifuge tube and dissolved in 1ml HPLC grade methanol and vertexed for 2-3 minutes. Graphitized carbon black (GCB) @ 0.2g was added into the solution and again vetexted for 1 minute for the homogenization of the mixture. The function of GCB was used to remove the pigmentations and sterols from the extracted solvents. The recovery rate was recorded maximum when 0.2 g of GCB were used [125]. The mixture was centrifuged for 5 minutes at 5000 revolutions per min at 27°C; however this procedure was adopted twice to obtain good results. The transparent supernatant solvent were collected and evaporated to dryness with a gentle stream of nitrogen and then dissolved in 1mL methanol for further analysis [125].

### 6.6. Gas Chromatography-Mass Spectrophotometery (GC-MS) Analysis

GC-MS analysis was assessed on an Agilent 6890-5973N USA with the gas chromatograph set with an HP1 capillary column with model number was TG-5MS (30 m×250 um × 0.25 um) polydimethylsiloxane interfaced with Hewlett Packard (5973N) mass selective detector. The initial temperature was 70°C (2 min) and last temperature increased to 200°C with final time 10°C min^-1^ however the inlet temperature was set out 250°C having split ratio 10:1. MS quadruple and thermal aux temperatures were 285°C respectively. The MS scan range was 35-520 units and helium gas used as carrier with 1.0 mL min^−1^ flow rate [115].

### 6.7. Compound Identifications

Different collected chemicals compounds were identified and verified on molecular mass, molecular formula basis [126,127]. The comparative yield of compounds raw data was calculated based on gas chromatography (GC) areas with a FID correction factor which is specific, linear, sensitive, precise and accurate [115,128] method of measurement.

### 6.8. Statistical analyses

The accuracy of natural enemy pressure was estimated with latitude. One-way analysis of variance conducted to test the effect of latitude on different growth parameters, natural enemy’s abundance and phyto-chemical yield extractions [99]. Principal component and redundant analysis were conducted to explore the multivariate relationship between natural enemy’s abundance and different abiotic trends. This procedure was done to reduce the dimensionality in plant growth variables [99]. The achievement of the resultant Principal Component Analysis and RDA (axes 1 and 2 and or 3) were narrated as representative variables, that were linear mixture of the species development variables [57]. However, a generalized linear model (GLM) for multivariate analysis (MANOVA) was conducted to determine the significance of fixed effects using the type III sum of the square [61]. To investigate the variation among different sites, temperature and plant growth parameters during a field survey at eight different locations, nested ANOVAs were calculated. The analysis was conducted while considering environmental factors and plant height, diameter to develop a trend of abiotic factors with insect and pathogen abundance [53,57]. Moreover, principal component analysis (PCA) was conducted using R version 3.3.1 with “vegan” package used for RDA [99,129]. SPSS 13 was used for calculating generalized linear models (GLM’s), and the interaction of growth parameter and ecological factors with environmental factors were designed on Minitab 19 statistical software. The geographic coordinates of all the survey sites were recorded using GPS navigator (GPS-Hollox, Taiwan), and ArcGIS version 10.2 software (ESRI) was used to map the distribution of *Xanthium strumarium* at different latitudes [130].

## Acknowledgement

The survey conducted with the support of National Key R&D Program of China (2017YFC1200101), the National Natural Science Foundation of China (31470575, 31670545, and 31971557).

## Author Contributions

The following statements stated that “conceptualization, Y.L.F. and M.F.I.; methodology, Y.L.F.; software, M.F.I.; validation, Y.L.F., M.F.I.; formal analysis, M.F.I.; investigation, M.F.I.; resources, Y.L.F.; data curation, M.F.I.; writing—original draft preparation, M.F.I., Y.L.F.; writing—review and editing, M.F.I; Y.L.F.; visualization, M.F.I.; supervision, Y.L.F.; project administration, Y.L.F.; funding acquisition, Y.L.F.”,

